# *Mycobacterium tuberculosis* DprE1 inhibitor OPC-167832 is active against *Mycobacterium abscessus in vitro*

**DOI:** 10.1101/2022.09.14.508059

**Authors:** Jickky Palmae Sarathy, Matthew D. Zimmerman, Martin Gengenbacher, Véronique Dartois, Thomas Dick

## Abstract

The anti-tuberculosis candidate OPC-167832, an inhibitor of DprE1, was active against *Mycobacterium abscessus*. Resistance mapped to *M. abscessus dprE1*, suggesting target retention. OPC-167832 was bactericidal and did not antagonize activity of clinical anti-*M. abscessus* antibiotics. Due to its moderate potency compared to *Mycobacterium tuberculosis*, the compound lacked efficacy in a mouse model and is thus not a repurposing candidate. These results identify OPC-167832 – DprE1 as a lead-target couple for a *M. abscessus*-specific optimization program.

## MAIN TEXT

The non-tuberculous mycobacterium (NTM) and opportunistic pathogen *M. abscessus* can establish extremely difficult to treat lung infections (1–3). Complex antibiotic regimens, typically consisting a macrolide (clarithromycin or azithromycin), the aminoglycoside amikacin, and a β-lactam (cefoxitin or imipenem), are administered often for years and show low cure rates (4–8). In brief, there is no reliable cure against *M. abscessus* lung disease. The *M. abscessus* drug pipeline is thinly populated (9), and new repurposing candidates and lead-target couples are sorely needed (10).

Decaprenylphosphoryl-β-D-ribose oxidase (DprE1) has emerged as an attractive target for anti-tuberculosis (TB) drug development (11–13). The enzyme, catalyzing the formation of decaprenyl-phospho-arabinose (DPA), is essential for growth and viability of *Mycobacterium tuberculosis* (14–16). DPA serves as a precursor for the synthesis of arabinogalactan, a critical component of the mycobacterial cell wall (14). Inhibitors of *M. tuberculosis* DprE1 have been identified from various structural scaffolds and show potent activity *in vitro* and in mouse models (13, 17, 18). BTZ-043, PBTZ-169, OPC-167832 and TBA-7371 have progressed to phase I or II clinical trials for TB (19).

Transposon mutagenesis studies have shown that *M. abscessus dprE1* (*mab_0192c*) is genetically essential (20). Whether *M. abscessus* DprE1 is a vulnerable target whose inhibition would translate into whole-cell antimicrobial activity has not been established. BTZ-043 and its analog PBTZ-169 have been tested for activity against *M. abscessus* and both were found to be inactive (21, 22). This is likely due to an amino acid polymorphism in *M. abscessus* DprE1. BTZ-043 and PBTZ-169 form covalent adducts with cysteine 387 in *M. tuberculosis* DprE1 as their on-target mechanism of action (22–24). *M. abscesuss* DprE1 displays alanine at the corresponding amino acid residue position, thus preventing covalent adduct formation and enzyme inhibition by the covalent inhibitors (22–24).

Here, we tested the growth inhibitory activity of the non-covalent DprE1 inhibitors OPC-167832 and TBA-7371 (25, 26). The minimum inhibitory concentration (MIC) of the compounds against the type-strain *M. abscessus* subsp. *abscessus* ATCC 19977 (American Type Culture Collection) was determined in Middlebrook 7H9 broth (BD) using the broth microdilution method with optical density at 600 nm (OD_600_) as the readout, as described previously (27). MIC was defined as 90% growth inhibition compared to the drug-free culture. While TBA-7371 (MedChem Express) was inactive (MIC > 100 μM), the dihydrocarbostyril OPC-167832 (MedChem Express) was found to be active (MIC = 5.2 μM). To determine whether the activity of OPC-167832 against the type-strain was retained against the broader *M. abscessus* complex (28), MICs were measured against the reference strains of the two other subspecies, *M. abscessus* subsp. *bolletii* CCUG 50184T and *M. abscessus* subsp. *massiliense* CCUG 48898T (Culture Collection University of Goteborg), and against a panel of clinical isolates which include *erm41*-harboring macrolide-resistant strains (29, 30). Potency was largely consistent across the *M. abscessus* complex with MICs ranging from 5.2 to 15 μM (Table 1).

**TABLE 1.** MIC of OPC-167832 against *M. abscessus* complex

| *M. abscessus* strain | *erm41*<br>*sequevar*<sup>b</sup> | Clarithromycin<br>susceptibility | MIC (μM)<sup>c</sup> |  |
| --- | --- | --- | --- | --- |
|  |  |  | OPC-167832 | Clarithromycin |
| Reference strains |  |  |  |  |
| subsp. *abscessus* ATCC 19977 | T28 | Resistant | 5.2 | 1.2 |
| subsp. *bolletii* CCUG 50184-T | T28 | Resistant | 5.4 | 3.2 |
| subsp. *massiliense* CCUG 48898-T | Deletion | Sensitive | 6 | 0.6 |
| Clinical isolates<sup>a</sup> |  |  |  |  |
| subsp. *abscessus* Bamboo | C28 | Sensitive | 10 | 0.8 |
| subsp. *abscessus* K21 | C28 | Sensitive | 15 | 2 |
| subsp. *abscessus* M9 | T28 | Resistant | 10 | 5.4 |
| subsp. *abscessus* M199 | T28 | Resistant | 10 | 4.8 |
| subsp. *abscessus* M337 | T28 | Resistant | 7 | 2.9 |

**TABLE 1** MIC of OPC-167832 against *M. abscessus* complex
|  |  |  |  |  |
| --- | --- | --- | --- | --- |
| subsp. *abscessus* M404 | C28 | Sensitive | 8.5 | 0.8 |
| subsp. *abscessus* M421 | T28 | Resistant | 9.3 | 1.3 |
| subsp. *bolletii* M232 | T28 | Resistant | 8.8 | 5.4 |
| subsp. *bolletii* M506 | C28 | Sensitive | 6 | 0.6 |
| subsp. *massiliense* M111 | Deletion | Sensitive | 11 | 0.7 |

<sup>a</sup> *M. abscessus* Bamboo, *M. abscessus* K21 and the other clinical isolates were characterized and reported previously (32, 39, 40).
<sup>b</sup> *erm41* is the methylase gene responsible for inducible clarithromycin resistance. The “C28” and “deletion” sequevars are inactive *erm41* alleles and result in susceptibility to clarithromycin, while the “T28” sequevar is functional and confers inducible resistance against clarithromycin (29, 30).
<sup>c</sup> MIC determination was carried out three times independently and the results are presented as mean values. Clarithromycin (Sigma-Aldrich) was used as assay control (34).

The μM activity against the NTM is in stark contrast to the nM activities of OPC-167832 reported for *M. tuberculosis* (25). The dramatic *in vitro* potency difference of the TB drug candidate against *M. abscessus* suggests that OPC-167832 is likely not a repurposing candidate for the treatment of this lung disease. This was confirmed by *in vivo* pharmacokinetic – pharmacodynamic analyses. The plasma concentration-time profile upon oral administration of OPC-167832 in uninfected CD-1 mice (Charles River Laboratories) was determined by measuring the plasma concentrations of the compound via high pressure liquid chromatography coupled to tandem mass spectrometry (LC-MS/MS), as described previously (31). Dosing 20 or 100 mg/kg resulted in a plasma concentration versus time curve above the MIC for *M. tuberculosis* (25) for most of a 24 h interval, however the MIC for *M. abscessus* was not reached (Fig. 1A). As increasing the dose from 100 to 200 mg/kg did not result in a significant increase of exposure (Fig. 1A), 100 mg/kg was chosen as the highest dose for an efficacy study in a *M. abscessus* mouse model. NOD.CB17-Prkdc^scid^/NCrCrl mice (NOD SCID; Charles River Laboratories) were infected with *M. abscessus* K21 as described previously (32) and treated once daily for 10 days with orally administered OPC-167832 (50 or 100 mg/kg), the positive control clarithromycin (250 mg/kg) or drug-free vehicle. As expected, OPC-167832 treatment did not result in a statistically significant reduction of the lung bacterial burden (Fig. 1B). Plasma concentrations of OPC-167832 were measured 3 h and 24 h after the last dose, confirming similar concentrations in infected and naïve mice (Fig. 1A and 1C). Together, these *in vivo* analyses suggest that OPC-167832 is not a repurposing candidate for *M. abscessus* lung disease due to its moderate μM *in vitro* potency compared to its nM activity against *M. tuberculosis*. All experiments involving live animals were approved by the Institutional Animal Care and Use Committee of the Center for Discovery and Innovation, Hackensack Meridian Health.

**FIG 1.**
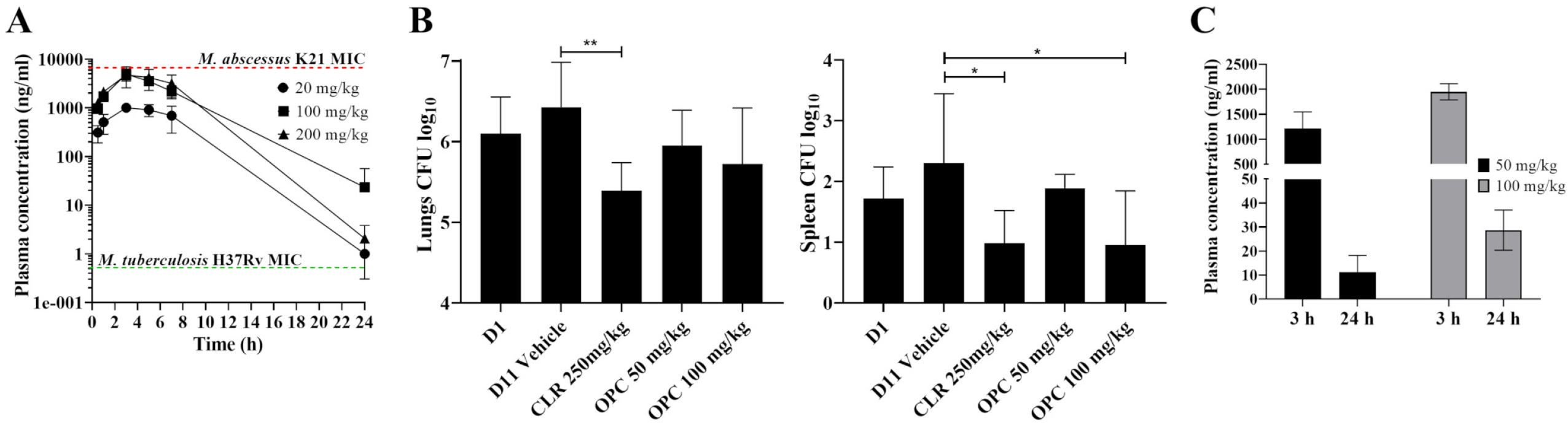
Pharmacokinetic profile and activity of OPC-167832 in mice. (A) Plasma concentration versus time profile of OPC-167832. Female CD-1 mice received a single dose of 20, 100 or 200 mg/kg of OPC-167832 formulated in 5 % (w/vol) gum arabic solution (Sigma-Aldrich) by oral gavage. Blood samples were collected from the tail vein at 0.5, 1, 3, 5, 7 and 24 h after drug administration and plasma concentration of OPC-167832 was measured by high pressure liquid chromatography coupled to tandem mass spectrometry (LC-MS/MS). The MIC of OPC-167832 against *M. abscessus* K21 (15 μM / 6852.6 ng/ml, Table 1) is indicated by the red dotted line. The reported MIC of OPC-167832 against *M. tuberculosis* H37Rv (1.1 nM / 0.5 ng/ml) is indicated by the green dotted line (25). (B) *In vivo* efficacy of OPC-167832 against *M. abscessus* in a NOD SCID mouse model. NOD SCID mice were infected intranasally with *M. abscessus* K21. Starting one day post-infection (D1), OPC-167832 (50 or 100 mg/kg, formulated in 5 % gum arabic solution), the positive control clarithromycin (250 mg/kg, formulated in 0.5% carboxymethyl cellulose–0.5% Tween 80–sterile water) or drug-free OPC-167832 vehicle were orally administered to infected mice for 10 consecutive days via oral gavage. 24 h after the last dose (11 days post-infection), all mice were euthanized, and their lungs and spleen were aseptically removed prior to homogenization. Serial dilutions of organ homogenates were plated onto Middlebrook 7H11 agar (BD) to quantify lung and spleen bacterial load on day 1 post infection (D1) and after administration of drug-free vehicle (D11 vehicle), OPC-167832 (OPC) and clarithromycin (CLR). Mean and standard deviation are shown for each treatment group (n=6). Statistical significance was determined using one-way analysis of variance (ANOVA) multi-comparison and Dunnett’s post-test (*, p < 0.05; **, p < 0.01). The experiment was carried out twice, showing similar results and one representative dataset is shown. (C) Plasma concentrations of OPC-167832 in infected NOD SCID mice 3 and 24 h after the last dose in the efficacy experiment shown in (B). The graphs were generated using GraphPad Prism 9 software.

To determine whether OPC-167832 retains DprE1 as its target in *M. abscessus* and inform future lead optimization efforts, spontaneous resistant mutants were isolated using the type-strain *M. abscessus* ATCC 19977 on Middlebrook 7H10 agar as described previously (31). 20 x 10^9^ CFU were plated on agar medium containing 16x MIC (MIC = 5.2 μM, Table 1), the lowest OPC-167832 concentration that suppressed growth of wild type colonies, resulting in a frequency of resistance of 10^-9^/CFU. The experiment was repeated with another independently grown culture, yielding a similar frequency of resistance. Twelve randomly selected OPC-167832-resistant strains (OPC_RM1 to OPC_RM12) from the two selection experiments achieved pronounced resistance with ~30 to 100-fold higher MIC than the parent strain (Table 2). Whole genome sequencing (Novogene Corporation Inc.), followed by Sanger sequencing (Genewiz Inc.), revealed that the 12 OPC-167832-resistant strains comprised two genotypic classes. Six strains (OPC_RM1 to OPC_RM6) harbored four different missense mutations in the *M. abscessus* homolog of *dprE1*, while the other six resistant strains (OPC_RM7 to OPC_RM12) harbored five different missense mutations in the homolog of *sigA* (*mab_3009*), which encodes the essential sigma factor A that assists the RNA polymerase in recognizing promoters of target genes (Table 2) (33).

**TABLE 2.**
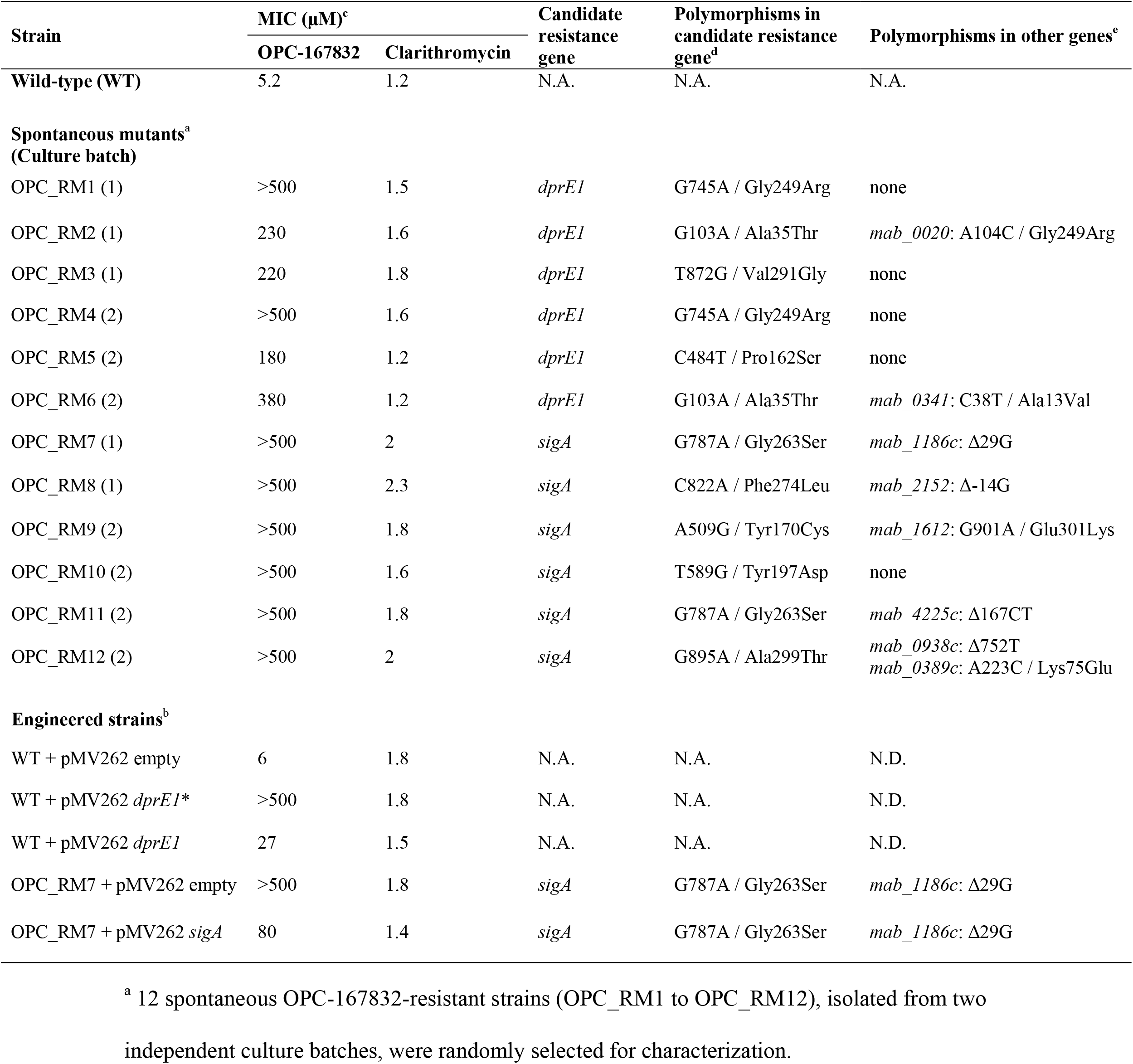
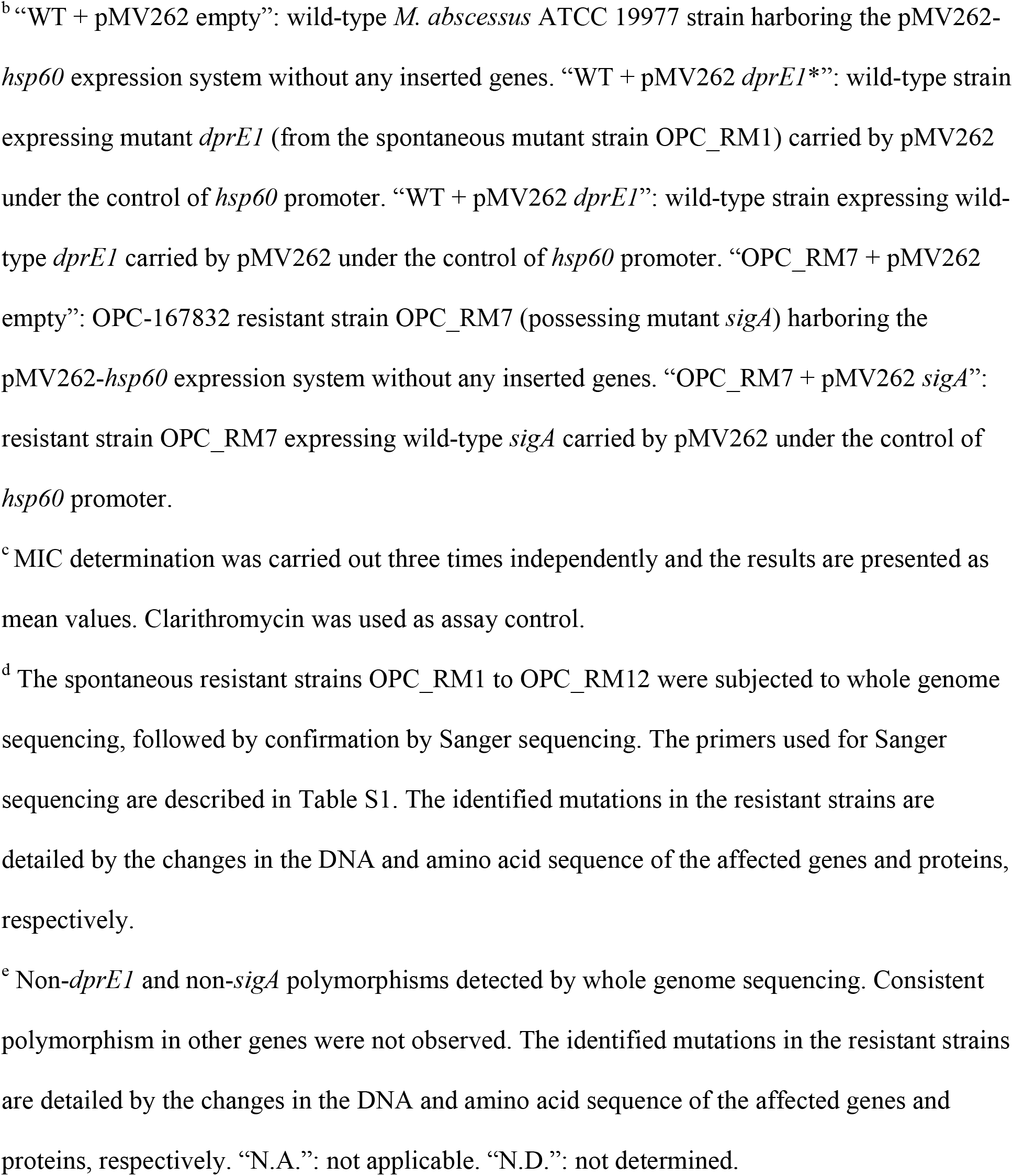
Characterization of spontaneous and engineered OPC-167832-resistant *M. abscessus* ATCC 19977 strains

| Strain | MIC (μM)<sup>c</sup> |  | Candidate resistance gene | Polymorphisms in candidate resistance gene<sup>d</sup> | Polymorphisms in other genes<sup>e</sup> |
| --- | --- | --- | --- | --- | --- |
|  | OPC-167832 | Clarithromycin |  |  |  |
| Wild-type (WT) | 5.2 | 1.2 | N.A. | N.A. | N.A. |
| **Spontaneous mutants<sup>a</sup>**<br>(Culture batch) |  |  |  |  |  |
| OPC_RM1 (1) | >500 | 1.5 | *dprE1* | G745A / Gly249Arg | none |
| OPC_RM2 (1) | 230 | 1.6 | *dprE1* | G103A / Ala35Thr | *mab_0020*: A104C / Gly249Arg |
| OPC_RM3 (1) | 220 | 1.8 | *dprE1* | T872G / Val291Gly | none |
| OPC_RM4 (2) | >500 | 1.6 | *dprE1* | G745A / Gly249Arg | none |
| OPC_RM5 (2) | 180 | 1.2 | *dprE1* | C484T / Pro162Ser | none |
| OPC_RM6 (2) | 380 | 1.2 | *dprE1* | G103A / Ala35Thr | *mab_0341*: C38T / Ala13Val |
| OPC_RM7 (1) | >500 | 2 | *sigA* | G787A / Gly263Ser | *mab_1186c*: Δ29G |
| OPC_RM8 (1) | >500 | 2.3 | *sigA* | C822A / Phe274Leu | *mab_2152*: Δ-14G |
| OPC_RM9 (2) | >500 | 1.8 | *sigA* | A509G / Tyr170Cys | *mab_1612*: G901A / Glu301Lys |
| OPC_RM10 (2) | >500 | 1.6 | *sigA* | T589G / Tyr197Asp | none |
| OPC_RM11 (2) | >500 | 1.8 | *sigA* | G787A / Gly263Ser | *mab_4225c*: Δ167CT |
| OPC_RM12 (2) | >500 | 2 | *sigA* | G895A / Ala299Thr | *mab_0938c*: Δ752T<br>*mab_0389c*: A223C / Lys75Glu |
| **Engineered strains<sup>b</sup>** |  |  |  |  |  |
| WT + pMV262 empty | 6 | 1.8 | N.A. | N.A. | N.D. |
| WT + pMV262 *dprE1* * | >500 | 1.8 | N.A. | N.A. | N.D. |
| WT + pMV262 *dprE1* | 27 | 1.5 | N.A. | N.A. | N.D. |
| OPC_RM7 + pMV262 empty | >500 | 1.8 | *sigA* | G787A / Gly263Ser | *mab_1186c*: Δ29G |
| OPC_RM7 + pMV262 *sigA* | 80 | 1.4 | *sigA* | G787A / Gly263Ser | *mab_1186c*: Δ29G |

<sup>a</sup> 12 spontaneous OPC-167832-resistant strains (OPC\_RM1 to OPC\_RM12), isolated from two
independent culture batches, were randomly selected for characterization.
<sup>b</sup>“WT + pMV262 empty”: wild-type *M. abscessus* ATCC 19977 strain harboring the pMV262- *hsp60* expression system without any inserted genes. “WT + pMV262 *dprE1*”\*: wild-type strain expressing mutant *dprE1* (from the spontaneous mutant strain OPC\_RM1) carried by pMV262 under the control of *hsp60* promoter. “WT + pMV262 *dprE1*”: wild-type strain expressing wild- type *dprE1* carried by pMV262 under the control of *hsp60* promoter. “OPC\_RM7 + pMV262 empty”: OPC-167832 resistant strain OPC\_RM7 (possessing mutant *sigA*) harboring the pMV262-*hsp60* expression system without any inserted genes. “OPC\_RM7 + pMV262 *sigA*”: resistant strain OPC\_RM7 expressing wild-type *sigA* carried by pMV262 under the control of *hsp60* promoter.
<sup>c</sup> MIC determination was carried out three times independently and the results are presented as mean values. Clarithromycin was used as assay control.
<sup>d</sup> The spontaneous resistant strains OPC\_RM1 to OPC\_RM12 were subjected to whole genome sequencing, followed by confirmation by Sanger sequencing. The primers used for Sanger sequencing are described in Table S1. The identified mutations in the resistant strains are detailed by the changes in the DNA and amino acid sequence of the affected genes and proteins, respectively.
<sup>e</sup> Non-*dprE1* and non-*sigA* polymorphisms detected by whole genome sequencing. Consistent polymorphism in other genes were not observed. The identified mutations in the resistant strains are detailed by the changes in the DNA and amino acid sequence of the affected genes and proteins, respectively. “N.A.”: not applicable. “N.D.”: not determined.

To confirm that the observed polymorphisms detected in *dprE1* and *sigA* are indeed responsible for resistance to OPC-167832, merodiploid strains were engineered using custom-synthesized pMV262-*hsp60*-based expression systems (Genewiz Inc.) as described previously (34). To confirm involvement of *dprE1* missense mutations, a copy of the mutant *dprE1* allele from a representative resistant strain (OPC_RM1, Table 2) was constitutively expressed under the control of the *hsp60* promoter in wild-type *M. abscessus* ATCC 19977. As expected, the strain expressing the mutant *dprE1* allele displayed high level resistance to OPC-167832 (Table 2). To exclude the possibility that the observed resistance was caused by mere overexpression of the *dprE1* gene, as opposed to the missense mutation harbored by the mutant *dprE1* allele, the wild-type allele of *dprE1* was expressed under the control of the *hsp60* promoter in wild-type *M. abscessus* ATCC 19977. This resulted in low-level resistance to OPC-167832 (Table 2), indicating that the missense mutations are the major contributors to the resistance phenotype. Together, these genetic analyses suggest that DprE1 is a target of OPC-167832 in *M. abscessus*. To confirm that the observed polymorphisms in *sigA* cause resistance to OPC-167832, one representative *sigA* mutant strain (OPC_RM7, Table 2) was complemented with a wild-type copy of the *sigA* gene that was constitutively expressed under the control of the *hsp60* promoter. Expression of wild-type *sigA* in the mutant background partially restored sensitivity to OPC-167832 (Table 2), suggesting that the missense mutations observed in *sigA* are the cause of resistance to OPC-167832. A few reports describe mutations in *sigA* causing drug resistance in other bacteria, apparently by reprogramming the transcriptome (35, 36). How mutations in *M. abscessus sigA* cause resistance against OPC-167832 remains to be determined.

To further evaluate the attractiveness of OPC-167832 – DprE1 as a lead-target couple, *in vitro* bactericidal activity and *in vitro* drug-drug potency interactions with anti-*M. abscessus* antibiotics were determined using *M. abscessus* ATCC 19977 as described previously (27). OPC-167832 was highly bactericidal, with a 3-log kill at 4x MIC (Fig. 2). Absence of antagonism with clarithromycin, amikacin (Sigma-Aldrich), cefoxitin (MedChem Express) or imipenem (Cayman Chemical) (Table 3), together with the clean drug-drug interaction profile of OPC-167832 as required under multi-drug TB therapy (25, 37, 38), suggest that dihydrocarbostyril analogs are compatible with the current standard of care for *M. abscessus* lung disease.

**FIG 2.**
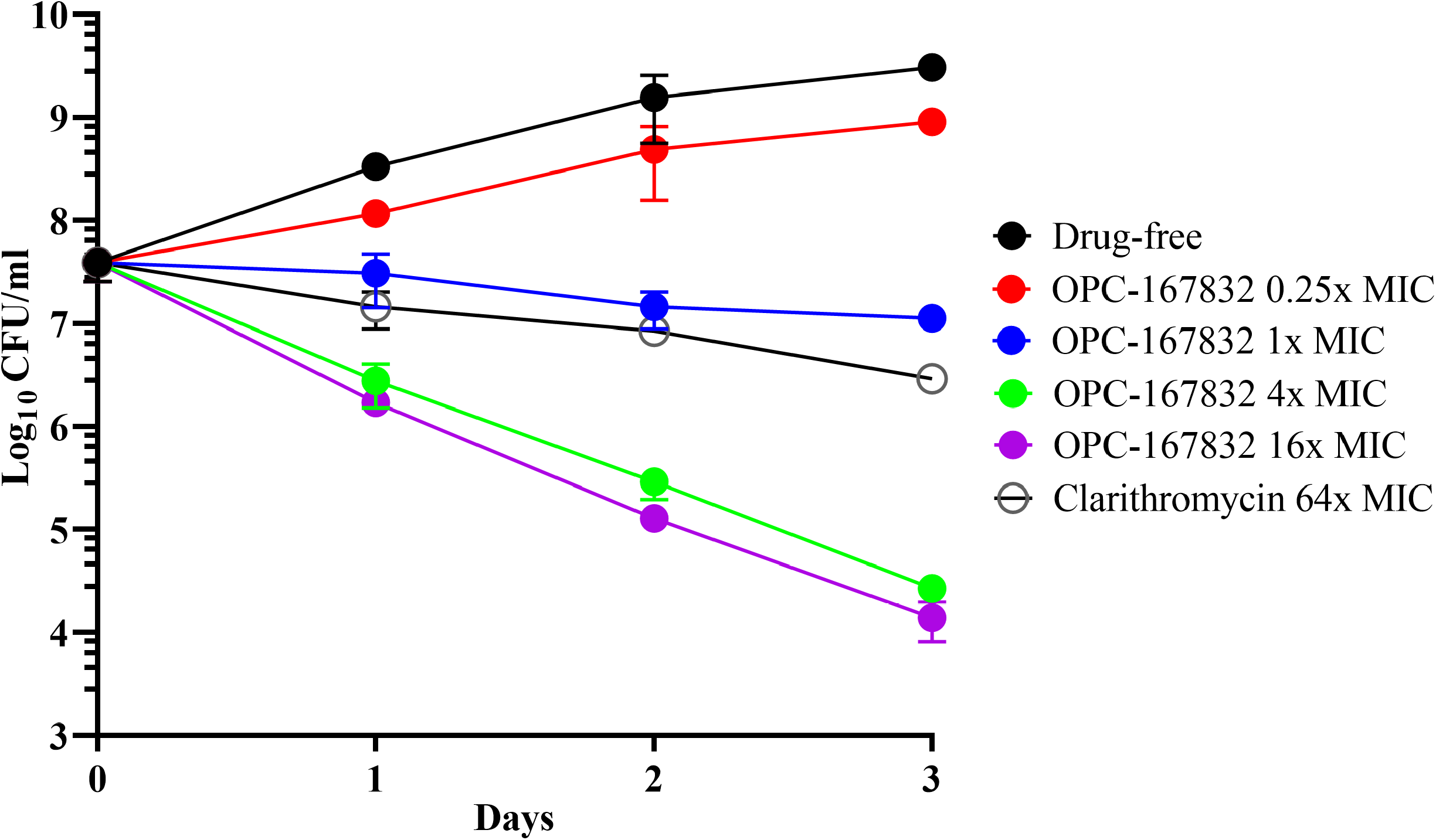
*In vitro* time-kill activity of OPC-167832 against *M. abscessus* ATCC 19977. *M. abscessus* cultures were grown in Middlebrook 7H9 and treated with 0.25x, 1x, 4x, and 16x MIC of OPC-167832 (MIC: 5.2 μM, Table 1) over a period of three days and CFU/ml were measured by plating samples on Middlebrook 7H10 agar. Clarithromycin was used as a negative control at 64x MIC (MIC: 1.2 μM, Table 1). The experiment was carried out three times independently and the results are presented as mean values with standard deviations displayed as error bars. The graph was generated using GraphPad Prism 9 software.

**TABLE 3.** *In vitro* drug-drug potency interaction between OPC-167832 and selected clinically used drugs against *M. abscessus* ATCC 19977

| Drugs<sup>a</sup> | Class | Target | MIC (μM)<sup>b</sup> |  | FICI<sup>c</sup> | Outcome<sup>c</sup> |
| --- | --- | --- | --- | --- | --- | --- |
|  |  |  | Alone | In combination |  |  |
| OPC-167832 | 3,4-dihydrocarbostyryl | DprE1 | 5.2 | 1.9 | 0.6 | Additivity |
| Clarithromycin | Macrolide | 50S ribosomal subunit | 1.2 | 0.3 |  |  |
| OPC-167832 | 3,4-dihydrocarbostyryl | DprE1 | 5.2 | 1.6 | 0.9 | Additivity |
| Amikacin | Aminoglycoside | 30S ribosomal subunit | 30 | 17 |  |  |
| OPC-167832 | 3,4-dihydrocarbostyryl | DprE1 | 5.2 | 1.7 | 0.6 | Additivity |
| Cefoxitin | β-lactam | Peptidoglycan biosynthesis transpeptidases | 26 | 7.8 |  |  |
| OPC-167832 | 3,4-dihydrocarbostyryl | DprE1 | 5.2 | 1.5 | 0.4 | Synergy |
| Imipenem | β-lactam | Peptidoglycan biosynthesis transpeptidases | 25 | 3.6 |  |  |

<sup>a</sup> To determine possible antagonisms between OPC-167832 and clinically used drugs, checkerboard analyses were carried using a 96-well plate format (41, 42). The effect of serially diluted OPC-167832 ranging from 0.39 μM to 25 μM was tested against the partner drugs ranging from 0.49 μM to 250 μM.
<sup>b</sup> MIC determination was carried out three times independently and the results are presented as mean values.
<sup>c</sup> The Fractional Inhibitory Concentration Index (FICI) was calculated as [(MIC of partner drug in combination / MIC of partner drug alone) + (MIC of OPC-167832 in combination / MIC of OPC-167832 alone)]. FICI ≤ 0.5 indicates synergy, FICI 0.5 – 4 indicates additivity (no interaction) and FICI > 4 indicates antagonism (43).

In conclusion, we identified OPC-167832 as the first whole-cell active inhibitor of *M. abscessus* DprE1, thus validating DprE1 as a vulnerable target in the opportunistic pathogen. The 1000-fold weaker activity of OPC-167832 against *M. abscessus* compared to *M. tuberculosis* results in unfavorable pharmacokinetic – pharmacodynamic parameters and lack of efficacy in a mouse model of *M. abscessus* infection. Thus, the TB drug candidate is unlikely to present a repurposing candidate for the treatment of *M. abscessus* lung disease. The reason for the pronounced potency difference against the two mycobacterial species remains to be determined and may involve target binding, uptake/excretion, or intra-bacterial metabolism (10). If the basis for the differential potency can be elucidated, OPC-167832 may present an attractive chemical starting point for a rational, pathogen-specific lead optimization program.

## ACKNOWLEDGMENTS

We are grateful to Wei Chang Huang (Taichung Veterans General Hospital, Taichung, Taiwan) for providing *M. abscessus* Bamboo, to Jeanette W.P. Teo (Department of Laboratory Medicine, National University Hospital, Singapore) for providing the collection of *M. abscessus* clinical M isolates, and to Sung Jae Shin (Department of Microbiology, Yonsei University College of Medicine, Seoul, South Korea) and Won-Jung Koh (Division of Pulmonary and Critical Care Medicine, Samsung Medical Center, Seoul, South Korea) for providing *M. abscessus* K21. Research reported in this work was supported by the National Institute of Allergy and Infectious Diseases of the National Institutes of Health under Award Number R01AI132374. The content is solely the responsibility of the authors and does not necessarily represent the official views of the National Institutes of Health.

## AUTHOR CONTRIBUTIONS

Investigation: J.P.S., M.D.Z., M.G.; Writing - Original Draft: J.P.S, T.D.; Writing - Review & Editing: all authors; Funding Acquisition: T.D.; Supervision: M.G., V.D., T.D.

## CONFLICT OF INTEREST STATEMENT

The authors declare no commercial or financial relationships that could be construed as a potential conflict of interest.

